# Barcoding and demultiplexing Oxford Nanopore native RNA sequencing reads with deep residual learning

**DOI:** 10.1101/864322

**Authors:** Martin A. Smith, Tansel Ersavas, James M. Ferguson, Huanle Liu, Morghan C Lucas, Oguzhan Begik, Lilly Bojarski, Kirston Barton, Eva Maria Novoa

## Abstract

Nanopore sequencing has enabled sequencing of native RNA molecules without conversion to cDNA, thus opening the gates to a new era for the unbiased study of RNA biology. However, a formal barcoding protocol for direct sequencing of native RNA molecules is currently lacking, limiting the efficient processing of multiple samples in the same flowcell. A major limitation for the development of barcoding protocols for direct RNA sequencing is the error rate introduced during the base-calling process, especially towards the 5’ and 3’ ends of reads, which complicates sequence-based barcode demultiplexing. Here, we propose a novel strategy to barcode and demultiplex direct RNA sequencing nanopore data, which does not rely on base-calling or additional library preparation steps. Specifically, custom DNA oligonucleotides are ligated to RNA transcripts during library preparation. Then, raw current signal corresponding to the DNA barcode is extracted and transformed into an array of pixels, which is used to determine the underlying barcode using a deep convolutional neural network classifier. Our method, *DeePlexiCon*, implements a 20-layer residual neural network model that can demultiplex 93% of the reads with 95.1% specificity, or 60% of reads with 99.9% specificity. The availability of an efficient and simple barcoding strategy for native RNA sequencing will enhance the use of direct RNA sequencing by making it more cost-effective to the entire community. Moreover, it will facilitate the applicability of direct RNA sequencing to samples where the RNA amounts are limited, such as patient-derived samples.

## INTRODUCTION

The appearance of third generation sequencing (TGS) technologies has revolutionized our ability to sequence genomes and transcriptomes (1, 2). In comparison to next-generation sequencing technologies, TGS have the ability to produce long sequencing reads, avoiding the hassle of fragmenting the RNA or DNA molecules into smaller pieces to then reassemble them back together. Furthermore, TGS technologies have the ability to sequence DNA and RNA without a PCR amplification step, thus allowing for direct detection of DNA and RNA modifications, with single nucleotide and single molecule resolution.

Direct sequencing of native RNA molecules (dRNAseq) can be achieved using the platform offered by Oxford Nanopore Technologies (ONT). This platform relies on the use of protein nanopores embedded in a lipidic membrane that are subjected to an electric field. Characteristic disruptions in electric current are measured as the charged molecule passes through the pore, enabling the observation of single molecules. Low translocation velocity of the RNA molecule is achieved through the association of motor proteins that regulate translocation of nucleic acid polymers, and the current intensity measurements can in turn be converted into sequence information using base-calling algorithms (3).

The first direct RNA sequencing protocol developed by ONT (SQK-RNA001) became commercially available in 2017 and was designed to sequence mRNAs (4), although later efforts have shown that this protocol can be adapted to sequence non-polyAed RNAs, such as ribosomal RNAs (5). The current ONT dRNAseq library preparation protocol comprises three main steps: (i) ligation of a double-stranded, pre-annealed DNA RT Adapter (RTA), which contains an oligo-dT overhang to anneal to poly(A)+ mRNAs; (ii) optional reverse transcription, which linearizes the RNA molecule into an RNA-DNA duplex; and (iii) ligation of the RNA sequencing adapter (RMX), which contains the motor protein that directs RNA molecules to the pores and regulates their translocation (**Figure 1A**). Currently, there are no manufacturer-provided protocols for molecular barcoding of direct RNA sequencing datasets, which would improve the cost-effectiveness of certain dRNAseq applications by combining multiple samples on the same consumable flow cell. Moreover, it would allow the use of dRNAseq in cases where the amount of RNA sample is limiting - current input RNA requirements for dRNASeq is 500 ng, which greatly limits the applicability of the technology.

**Figure 1:**
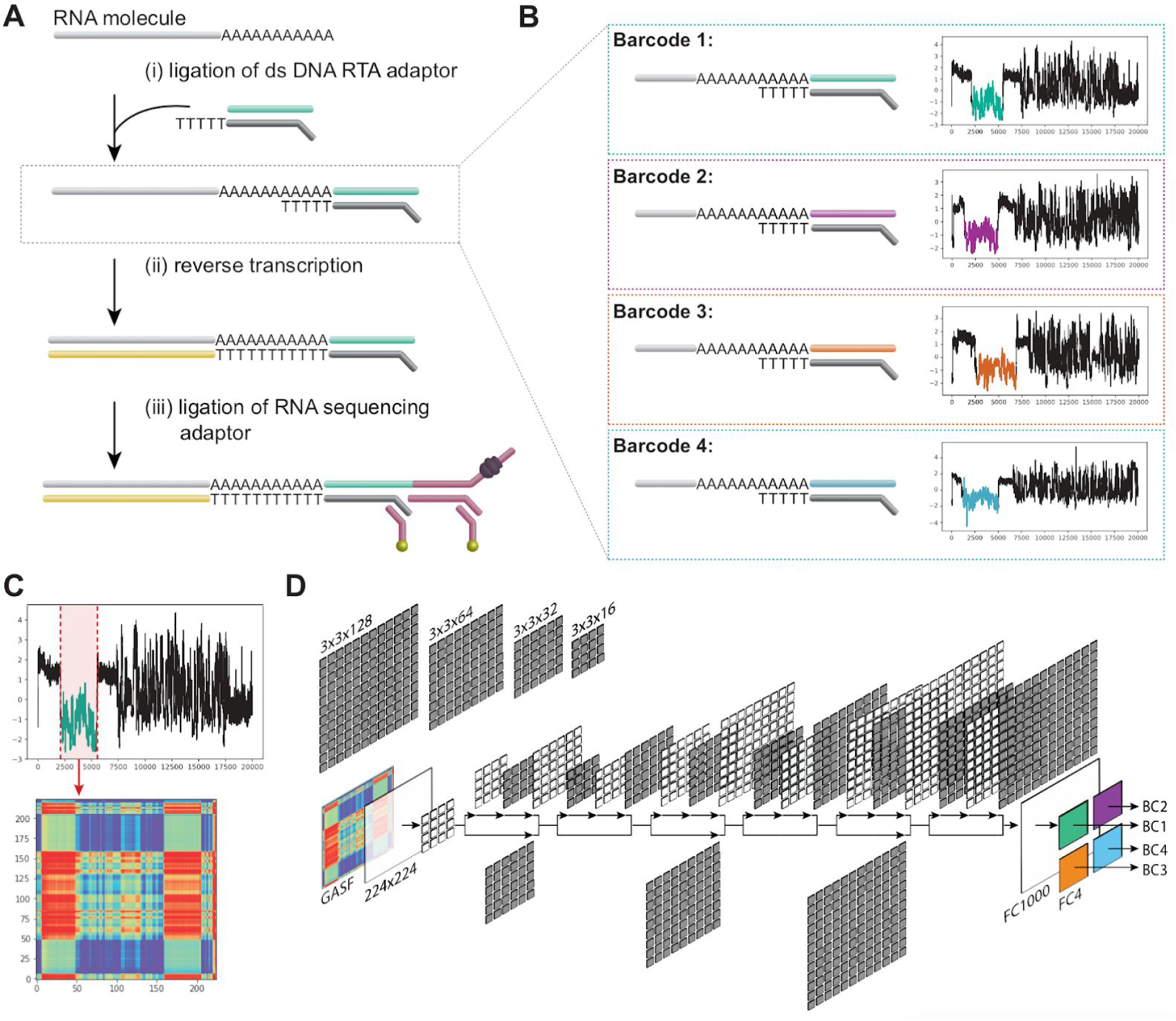
Direct RNA barcoding and demultiplexing. **(A)** Overview of Oxford Nanopore sample preparation protocol for native RNA sequencing. **(B)** Adaptation of (A) to include custom DNA barcodes. **(C)** Barcode segmentation and transformation, where the electric current associated with a barcode adapter (highlighted in red) is extracted and converted into an image using GASF transformation. **(D)** Deep learning is used to classify the segmented and GASF-transformed squiggle signals into their corresponding bins, without the need of base-calling the underlying sequence. The convolution architecture of the final residual neural network classifier (ResNet-20) described in this work: FC = Fully Connected layer.

Here we propose a novel strategy to barcode and efficiently demultiplex dRNAseq data (**Figure 1B**). Importantly, this strategy does not require additional ligation steps compared to the standard direct RNA sequencing library preparation, as it relies on the use of shuffled DNA oligonucleotides that are incorporated during the first ligation step. The DNA barcodes do not appear in the base-called fasta sequence --which are inferred from RNA-specific models--but their electronic signal is present in the raw sequencing data, which is used as input for our demultiplexing algorithm. Demultiplexing is performed via the transformation of raw FAST5 signals into images using Gramian Angular Summation Field (GASF), followed by classification using a deep residual neural network learning model (6). We demonstrate that our proposed methodology and algorithm is a highly effective strategy to multiplex direct RNA sequencing reads, yielding 99.9% specificity, while recovering 60% of the reads, -or 95.1% specificity with 93% of read recovery, if enhanced recovery is preferred-. The ability to barcode and accurately demultiplex direct RNA sequencing reads opens new avenues to enable nanopore native RNA sequencing of samples with limited RNA availability, such as patient-derived samples, as well as improves the cost-effectiveness of sequencing low diversity samples, such as target-enriched or *in-vitro* transcribed libraries.

## RESULTS

### Barcoding *in vitro* transcribed RNAs with shuffled DNA adapters

We designed three custom DNA barcode adapters by shuffling the double stranded sequence of the default ONT RTA adapter (**Figure 1B**). The three custom barcodes as well as the standard ONT RTA adapter were individually ligated to distinct *in vitro* transcribed RNA sequences (see *Methods* and **Table S1**). We performed five sequencing runs with the RTA and custom adapters: *replicates 1* and *3* contained four unique *Sequins* transcripts (7), while *replicates 2, 4*, and 5 contained four unique *Sequins* and four unique *Curlcake* sequences (8), with one of each ligated to a single adapter (**Table 1**). In addition, *replicate 3* was spiked-in with the manufacturer provided yeast *ENO2* control strand (RCS). Each run produced between 600,000-1,000,000 reads, which were basecalled and uniquely aligned to the reference sequences (**Table 1**, see also **Table S2**). The reference alignments were used to empirically demultiplex the sequences, thus establishing a truth set to train a barcode classifier.

**Table 1.**
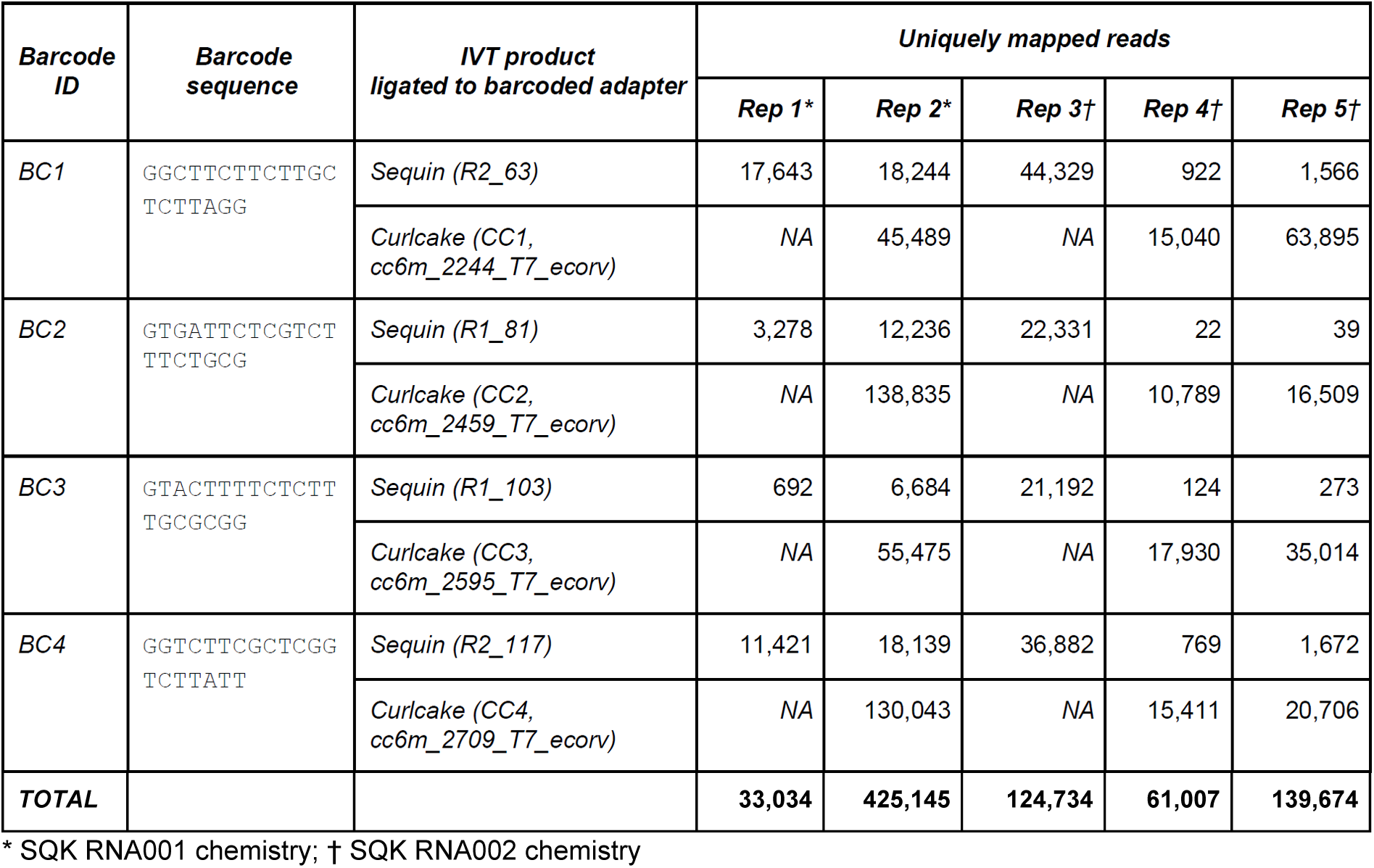
Mapping statistics from direct RNA sequencing runs.

### Extraction of barcode signals from raw FAST5 reads

Raw nanopore barcode signal data, consisting of a time series of electric current values, were extracted from the files corresponding to the uniquely mapped reads. Atomic structural differences between DNA and RNA produce conspicuously different mean current signal intensities, which can effectively be used to identify the boundaries of the proximal DNA adapter in the raw signal – a process henceforth referred to as *barcode segmentation*. We modified the *Segmenter* utility of *SquiggleKit* (9) *to create an automated workflow for barcode segmentation (termed B_roll*) that targets the lower average current level of the DNA barcodes by comparing the current of a given window to the average current of the read using a sliding window. We also tested a barcode segmentation strategy that uses raw current signal smoothing followed by convolutional transformation of the data (termed *B_conv*) to identify major current intensity change points along the read (see *Methods*). *B_roll* extracted signal from 74 out of 100 reads at an average speed of 0.013s per read, while *B_conv* extracted signal from 68/100 reads at an average speed of 2.45s per read (**Figure S1**). Although both methods proved sufficient for training a classifier (not shown), the *B_roll* method for barcode segmentation was chosen for subsequent analyses given its greater speed and recovery.

### Transformation of segmented barcode signals into 2D images

We reasoned that conveying raw current signal into a higher dimension could facilitate the recognition of similar patterns in the data by employing deep learning strategies for the downstream classification. Indeed, supervised machine learning using deep Convolutional Neural Networks (CNNs) and, in particular, deep Residual neural Networks (ResNet) have been shown to perform optimally for the classification of images (6, 10). To leverage the power of ResNet classifiers, we converted the raw signal corresponding to the extracted barcodes into an array of pixels using diverse image transformation strategies previously shown to be effective for subsequent CNN training and classification, including recurrence plots (RP) (11), Markov Transition Fields (MTF), Gramian Angular Difference Fields (GADF) and Gramian Angular Summation Fields (GASF) (12). An example of the different image transformations for a given raw signal segment can be found in **Figure S2**. GASF transformation was retained it was found to be substantially faster at computing images than the other methods (**Table 2**). Furthermore, the symmetrical images GASF produces generated slightly more accurate results than the non-symmetrical GADF images when tested in initial training experiments (not shown). **Figure 2** illustrates the conversion of segmented nanopore dRNAseq barcode signals into GASF images that were subsequently used for deep learning.

**Table 2.**
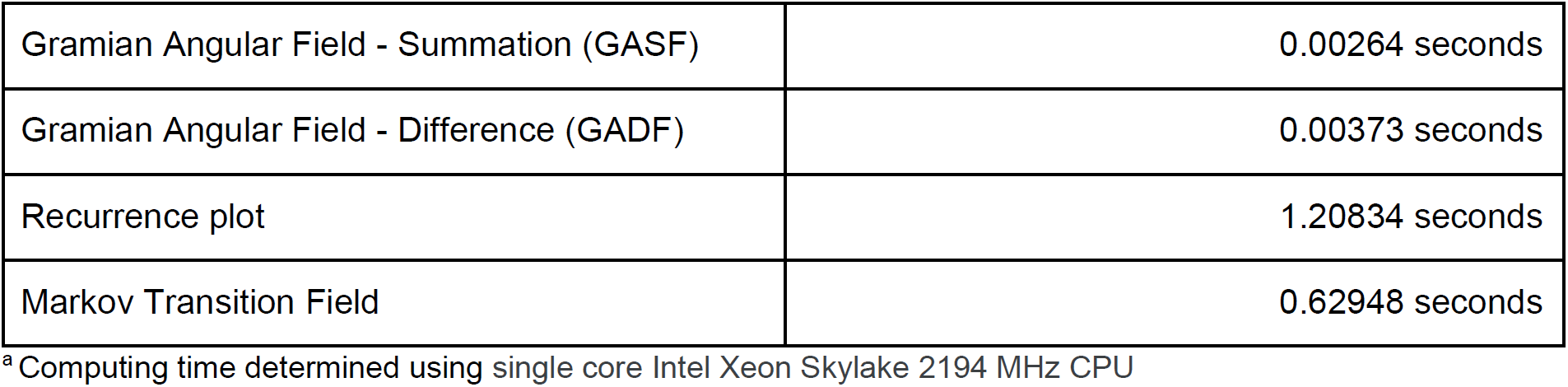
Average speed^a^ of signal to image conversions from 1000 runs.

|  |  |
| --- | --- |
| Gramian Angular Field - Summation (GASF) | 0.00264 seconds |
| Gramian Angular Field - Difference (GADF) | 0.00373 seconds |
| Recurrence plot | 1.20834 seconds |
| Markov Transition Field | 0.62948 seconds |

<sup>a</sup> Computing time determined using single core Intel Xeon Skylake 2194 MHz CPU

**Figure 2.**
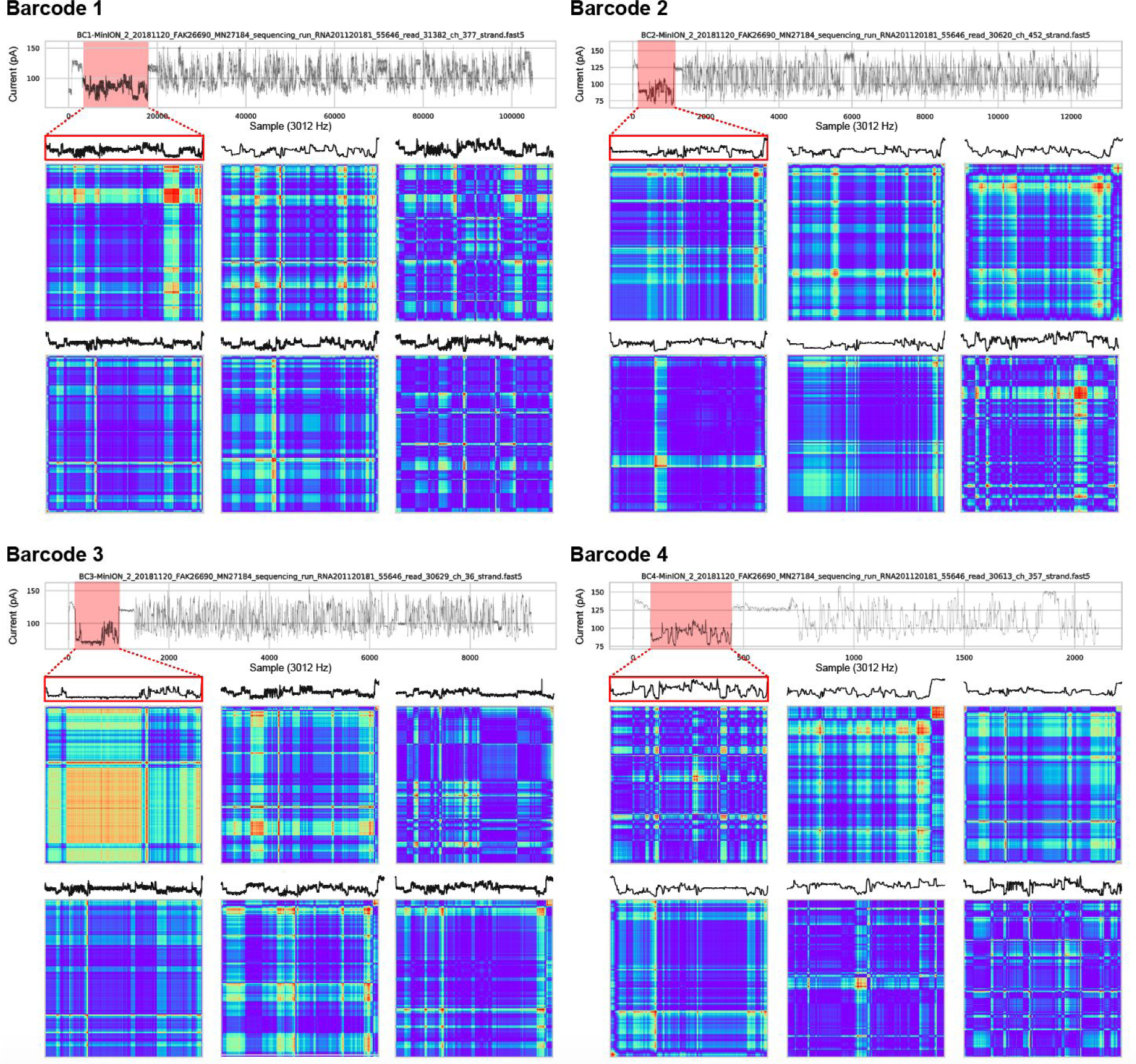
Barcode segmentation and signal transformation. A randomly selected example of barcode signal segmentation (red outline) for each of the four barcodes is shown with its corresponding GASF image below. An additional 5 randomly selected segmented barcode signals and their corresponding GASF images are shown for each of the four barcodes. Sequencing reads were drawn from replicate 2. GASF: Gramian Angular Summation Field.

### Deep residual networks to accurately classify raw signal barcodes

We combined sequencing data from replicates 2, 3 and 4 to train different CNN architectures using the GASF images generated from the segmented barcodes, which were previously disambiguated by aligning the base-called sequences of the ligated RNA sequenced to the reference sequence of their unique ligation templates. A total of 240k Images were divided into three groups of four barcodes for training, testing and validation at a ratio of 4:1:1, respectively (160K training : 40K testing : 40K withheld for validation). We compared a ResNet V2 implementation with 20 layers (ResNet-20, see **Figure 1D**) to a ResNet V2 with 56 layers (ResNet-56). We found that ResNet-20 was slightly better than ResNet-56 while being ⅓ smaller and three times faster (**Table 3**).

**Table 3.** Accuracy and training time of two residual neural networks on 4x Tesla V-100 GPUs

|  | ResNet-20 | ResNet-56 |
| --- | --- | --- |
| Training time | 6:21:52 | 19:21:26 |
| Accuracy/Loss<sup>a</sup> @ epoch 10 | 0.8956/0.3896 | 0.8825/0.4135 |
| Accuracy/Loss<sup>a</sup> @epoch 30 | 0.9735/0.1583 | 0.9356/0.2537 |
| Accuracy/Loss<sup>a</sup> @ epoch 45 | 0.9780/0.1448 | 0.9370/0.2489 |
| Training/Inference time per barcode | 3/3ms | 9/4ms |

<sup>a</sup>Accuracy and loss are calculated by the Keras framework (13), being (amount of correct guesses)/(total amount of guesses) and categorical cross entropy respectively.

The resulting ResNet-20 model was applied to the withheld validation set to assess its accuracy. Receiving Operator Characteristic (ROC) analysis revealed an Area Under the Curve of 0.998, a sensitivity of 98.9% and a false positive rate of 0.3% at maximal accuracy (99.4%) (**Table 4**, see also **Figure 3**), suggesting that the ResNet-20 model is highly tuned to the input and potentially overfitted, despite the latter being composed of three independent sequencing datasets.

**Table 4.** Accuracy and recovery of ResNet20 on the testing set, validation set, and two independent replicates.

| False positive rate (≤) | Deplexicon cutoff | Unclassified reads (%) | Accuracy (%) |
| --- | --- | --- | --- |
| *Testing Set (AUC=0.999)* |  |  |  |
| 0.01% | 1.0 | 69.9 | 85.3 |
| 0.1% | 0.9969 | 9.0 | 97.7 |
| 0.2%<sup>#</sup> | 0.8893<sup>#</sup> | 1.5<sup>#</sup> | 99.8<sup>#</sup> |
| 0.4%* | 0.0809* | 0.8* | 99.4* |
| 1.0% | 0.0139 | 0.7 | 99.1 |
| *Validation Set (AUC=0.998)* |  |  |  |
| 0.01% | 1.0 | 68.3 | 85.4 |
| 0.1% | 0.9991 | 17.2 | 95.6 |
| 0.3%# | 0.8164# | 1.1# | 99.4# |
| 0.4%* | 0.4396* | 1.4* | 99.6* |
| 1.0% | 0.0152 | 0.7 | 99.1 |
| *Independent Replicate (Rep. 1; AUC=0.954 )* |  |  |  |
| 0.01% | 1.0 | 97.5 | 75.6 |
| 0.1% | 1.0 | 86.1 | 78.4 |
| 1% | 0.9834 | 29.4 | 89.3 |
| 3.2%# | 0.7550# | 23.6# | 91.7# |
| 9.3%* | 0.1914* | 12.8* | 89.8 |
| *Independent Replicate (Rep. 5; AUC=0.987)* |  |  |  |
| 0.01% | 1 | 82.6 | 79.9 |
| 0.1% | 0.9983 | 39.6 | 89.1 |
| 1% | 0.8800 | 16.2 | 94.9 |
| 2.1%# | 0.6424# | 11.5# | 95.6# |
| 4.9%* | 0.2143* | 6.8* | 94.6* |

\* Optimal cutoff (Youden's J-statistic); # Maximum accuracy cutoff

**Figure 3.**
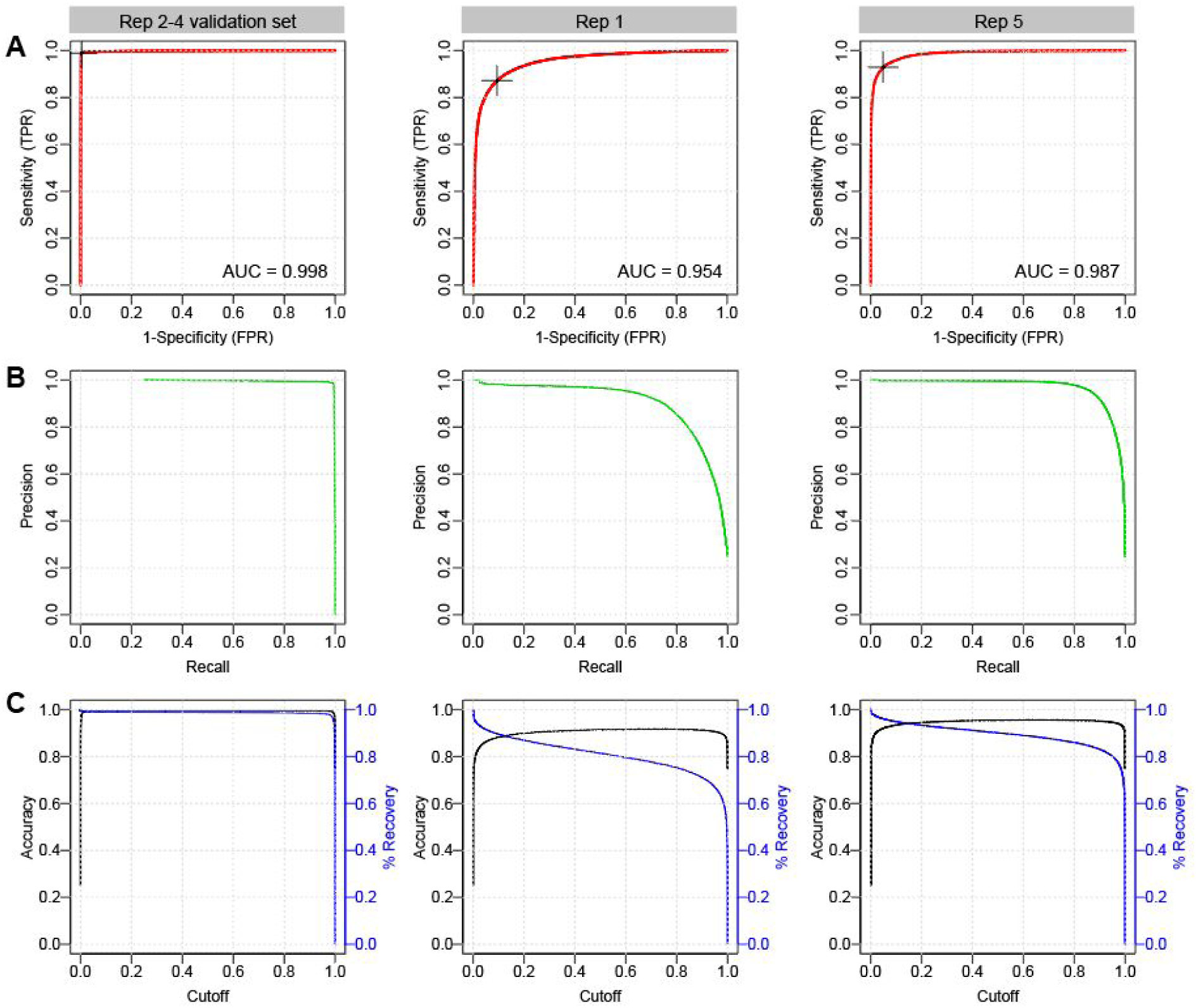
Performance of 2D convolutional neural network barcode classifier. **(A)** Receiving Operator Characteristic (ROC) analysis and Area Under the Curve (AUC) metrics of the final model on three evaluation sets: (i) Replicates 2-4 validation set (left column), which was generated from the same sequencing runs used to train the model, but were withheld from training; (ii) Replicate 1 set (middle column), composed of reads generated using the RNA001 library kit; and (iii) Replicate 5 set (right column), derived from an independent sequencing run using the RNA002 kit. Optimal Youden index (J statistic) is marked as a black cross on the ROC curve. **(B)** Accuracy (black) and percentage of reads recovered (blue) in function of the scoring threshold (cutoff) emitted by the trained model, for three different datasets presented in (A). **(C)** The associated precision recall curves on the 3 test sets.

To further evaluate the model’s accuracy and assess potential overfitting, we applied the model to two independent biological replicates (**Table 4**). The global accuracy of demultiplexing was slightly lower than the other replicates, with AUC values of 0.954 and 0.987 for rep. 1 and rep. 5, respectively (**Figure 3**). These slightly lower AUC values suggest that the ResNet-20 model may indeed be slightly overfitted to the sequencing data used for training, but nonetheless remains highly accurate at classifying reads from independent sequencing runs generated with different chemistries (RNA001 and RNA002, see Discussion).

## DISCUSSION

In the last decade, third generation sequencing technologies (TGS) have emerged as powerful methods to comprehensively study the (epi)transcriptome (14). In contrast to next generation sequencing technologies, TGS are not limited by read length, and consequently, do not require prior fragmentation of the RNA or cDNA molecules, providing transcriptome-wide maps of full-length molecules.

In 2017, the direct RNA sequencing (dRNAseq) technology appeared, making it possible for the first time to sequence native RNA molecules. Importantly, this technology could also identify chemical RNA modifications present in the native RNA molecules (4, 5, 8, 15), as well as estimations for their polyA-tail lengths (16, 17). However, a major caveat of dRNAseq is the amount of poly(A)-selected RNA material that is needed, i.e., typically 500ng of poly(A)+ RNA. Unfortunately, such amounts are difficult to obtain from biological samples, greatly limiting the applicability of this technology. In this regard, multiplexing samples in the same flow cell would allow this technology to be applied in situations where the amount of input RNA is limiting, as well as decrease the sequencing cost per sample. Unfortunately, ONT does not currently offer the possibility of multiplexing dRNAseq libraries.

In contrast to dRNAseq libraries, ONT does offer barcoding strategies for cDNA libraries, which rely on direct ligation of DNA adapters to the cDNA sequences. In this scenario, both the barcode and the cDNA sequence can be easily base-called under a DNA model. However, this is not possible in the context of RNA reads, as the adapter is DNA, and therefore, cannot be properly base-called under an RNA model. Alternatively, one could base-call the DNA adapter using the DNA model; however, this is not possible because the translocation speed of RNA reads (70bp/s) differs from that of DNA reads (450bp/s)

Here, we propose a novel strategy that relies on the use of deep Neural Networks (DNNs) to demultiplex dRNAseq libraries without the need of base-calling. Specifically, our strategy relies on conversion of the barcoded DNA adapter region into images, which are fed onto the trained DNNs to determine the underlying barcode (**Figure 1**). DNNs have been widely used in signal and time-series analysis problems, including speech recognition and electrical and optical signal coding-decoding (18). Compounding this fact, many of the recently developed DNA base-callers for nanopore signals rely on the use of DNNs, such as DeepNano (19), DeepSignal (20) or Chiron (21). Similarly, previous efforts have shown that nanopore DNA barcodes can be correctly classified using 1D Convolutional Neural Networks (CNNs) (22). Here we employ 2D CNNs, which are widely used in computer vision and pattern recognition (23), for direct classification of raw current intensity signals. Using this strategy, we correctly classified 84% of reads at 99% specificity (**Table 3**), which corresponds to 96.5% precision (positive predictive value) and 94.9% accuracy. The performance of *DeePlexiCon* is superior to the standard sequence-based strategies employed for DNA multiplexing and is comparable to the signal-based DNA demultiplexing algorithm DeepBinner, which displays slightly higher sensitivity and precision (92% and 98.5%, respectively) (22). This is most likely due to the longer barcodes employed in DNA multiplexing compared to dRNAseq (40 nt vs 20 nt, respectively), providing twice as much discriminative information. Albeit not the focus of this manuscript, the signal transformation and use of 2D CNNs for barcode demultiplexing would also be suitable for nanopore sequencing of DNA molecules, which might offer an alternative to DeepBinner that employs a 1D-based deep learning barcode classifier. Future efforts can increase the number of barcodes to allow multiplexing of additional samples in the same flow cell.

We should note that in the library preparation of replicate 1, which was one of the two datasets used for independent validation of the demultiplexing accuracy (**Table 4**), the 4 barcoded samples were pooled after the first ligation step but prior to reverse transcription and clean up, which may lead to spurious ligation events. Moreover, this library was loaded onto a R9.5 flowcell, which bears a modified nanopore protein optimised for rapid adapter uptake, whereas the remaining replicates were loaded onto R9.4 flowcells (**Table S2**). Although observed sporadically in other sequencing runs, replicate 1 revealed an increased frequency of spurious (equal barcode assignment probabilities), chimera (multi-mapping reads) and dual barcode ligations (false-false positive assignments evidenced by visual and algorithmic confirmation of dual barcodes in the raw signal), which likely explains the lower–yet reasonable–accuracy for this sample (**Figure S4**). The presence of multiple barcodes in a read might occur due to free floating adapters in solution in conjunction with minimal time between the first adapter/barcode passage, and the next, with a true read attached. However, this may also be due to the lack of clear open pore signal, causing MinKNOW to miss the segmentation, and thus produce a single fast5 file with both events included. Nonetheless, *DeePlexiCon* was able to demultiplex the sample with respectable accuracy (92-96%), demonstrating the power of deep learning for disentangling noisy data.

## METHODS

### Synthetic sequences

‘Curlcake’ sequences (8) were ordered from General Biosystems. Curlcake plasmids were double digested overnight with EcoRV-BamHI-HF. Sequin plasmid constructs (R2_117_1, R2_63_3, R1_103_1 and R1_81_2), used commercially for RNA sequencing experiments as a spike-in control (7), were a kind gift from Dr. Tim Mercer (https://www.sequinstandards.com/). ‘Sequin’ plasmids were digested O/N with EcoRI-HF. After digestion, DNA was extracted with Phenol-Chloroform followed by ethanol precipitation. Plasmid digestion was confirmed by agarose gel (**Figure S3A**). Digestion product quality was assessed with Nanodrop before proceeding to *in vitro* transcription.

### *In vitro* transcription, capping and polyadenylation

Using 1 µg of purified digestion product as starting material, Curlcake *in vitro* transcribed (IVT) sequences were produced using the Ampliscribe™ T7-Flash™ Transcription Kit (Lucigen-ASF3507). Sequin IVT sequences were produced using SP6 Polymerase (NEB-M0207S), following the manufacturer’s recommendations. Each IVT reaction was incubated for 4 hours at 42 °C for Curlcake sequences and at 40 °C for Sequin sequences. *In vitro* transcribed RNA was then incubated with Turbo DNAse (Lucigen) for 15 minutes, followed by purification using the RNeasy Mini Kit (Qiagen-74104). Correct IVT product lengths for Sequins were confirmed using Bioanalyzer (**Figure S3B**). Each IVT product was 5’ capped using Vaccinia Capping Enzyme (NEB-M2080S) following the manufacturer’s recommendations. The capping reaction was incubated for 30 minutes at 37 °C. Capped IVT products were purified using RNA Clean XP Beads (Beckman Coulter-A66514). Curlcake IVT products were Poly(A)-tailed using the *E. coli* Poly(A) Polymerase kit (NEB-M0276S), following the manufacturer’s recommendations. Poly(A)-tailed RNAs were purified using RNA Clean XP beads. Correct IVT product lengths for Curlcakes were confirmed using TapeStation (**Figure S3C**). Concentration of IVT products was determined using Qubit Fluorometric Quantitation and purity was measured with NanoDrop 2000 Spectrophotometer (**Table S3**)

### Direct RNA library preparation and sequencing

Custom RT adaptors (IDT) were annealed in following conditions. Oligo A and B were mixed in annealing buffer (0.01M Tris-Cl pH7.5, 0.05M NaCl) to the final concentration of 1.4 uM each in a total volume of 75 µl. The mixture was incubated at 94 °C for 5 minutes and slowly cooled down (−0.1 °C/s) to room temperature. RNA library for direct RNA Sequencing (SQK-RNA001 for replicates 1 and 2; SQK-RNA002 for replicates 3, 4 and 5) was prepared following the ONT Direct RNA Sequencing protocol (Version DRS_9026_v1_revP_15Dec2016 for replicates 1 and 2; DRS_9080_v2_revI_14Aug2019 for replicates 3, 4 and 5).

For replicates 2, 3, 4 and 5, 500 ng total of each IVT product (4 *Curlcakes* and/or 4 *Sequins*, as described in **Table 1**) were individually ligated to pre-annealed custom RT adaptors (IDT) (**Table S2**) in four separate eppendorfs, using concentrated T4 DNA Ligase (NEB-M0202T), and were reverse transcribed using SuperScript III Reverse Transcriptase (Thermo Fisher Scientific-18080044). The products were purified using 1.8X Agencourt RNAClean XP beads (Fisher Scientific-NC0068576), washing with 70% freshly prepared ethanol. In total, 50 ng of reverse transcribed RNA from each reaction was pooled, and RNA Adapter (RMX), composed of sequencing adapters with motor protein, was ligated onto the RNA:DNA hybrid. The mix was purified using 1X Agencourt RNAClean XP beads, washing with Wash Buffer twice. The sample was then eluted in Elution Buffer and mixed with RNA Running Buffer prior to loading onto a primed R9.4.1 flow cell (replicates 2,3,4 and 5) or R9.5 flow cell (replicate 1), and ran on either a GridION (replicates 1 and 3) or MinION (replicates 2, 4 and 5) sequencer for 48h or less (until all pores were inactive).

For replicate 1, library preparation steps were mainly performed as described above, but with slight variations. Specifically, the pooling of barcoded samples was performed after ligation step with pre-annealed custom RT adaptors, prior to reverse transcription. This strategy was discarded for the subsequent replicates as we considered that there could be potential cross-ligation of barcodes and IVT products if the pooling was performed prior to clean up.

### Basecalling, mapping and organization of sequencing data

Reads were basecalled with Guppy version 3.1.5 on a GPU-enabled Sun Grid Engine high performance computing server (parameters “--chunks_per_runner 1500 --gpu_runners_per_device 1 --cpu_threads_per_caller 4 -x “cuda:0 cuda:1 cuda:2 cuda:3” -r” and configuration “rna_r9.4.1_70bps_hac.cfg”. Base called reads (fastq) were aligned to Sequin transcripts (R2_117_1, R2_63_3, R1_103_1 and R1_81_2) (7) in replicate 1, and to both Sequin and ‘Curlcake’ constructs (CC1, CC2, CC3 and CC4) in replicate 2, using minimap2 (24) with v.2.17-r943-dirty with parameters “-k 14 --secondary=no”. Reference fasta sequences used to map both *Sequin* and *Curlcake* reads can be found in **Table S4**. Mapped reads were filtered for unique targets and mapping quality (MAPQ==60), quantified and binned into four groups based on the ligated sequence against which they mapped to, and the associated raw signal data was extracted using the fast5_fetcher and SquigglePull modules from the SquiggleKit package (9). The resulting tab delimited files were used as input for barcode segmentation, i.e., identifying and extruding the signal associated with DNA adapter barcodes.

### Extraction (segmentation) of raw signal associate with barcodes

Barcode segmentation from raw signal was performed using two strategies. The first strategy, which we term *B_roll*, calculates the global mean of the signal over a rolling window (2000 signal points) and identifies DNA barcode edges by setting a threshold of the mean, relative to the standard deviation. This strategy was performed by running the dRNA_segmenter.py script from SquiggleKit, with default parameters (9). The second strategy, which we term *B_conv*, consisted in applying the discrete convolution operation of the numpy python package (25) to smooth the unidimensional signal data and manifest large shifts in the data, which facilitates the identification of boundaries delimiting the different sections of the sequencing read. The 2nd derivative of convolved signal was calculated using a rolling window of 1001 points by applying the Savitzky-Golay filter (26). Maximal absolute values of derivatives were considered as the most likely location of boundary signal points, i.e., adapter start and end points. Mean and standard deviation of the current intensities were considered to further refine the boundaries. The raw signal comprised between the two boundary points, identified by either strategy, was used as input for the following steps. The efficiency and accuracy of both methods was assessed by visually inspecting 100 start and stop sites in the segmentation output of both methods. It was found that while *B_conv* had a bias in setting the end-point boundary too early, thus reducing the number of viable full segments (**Figure S1**)

### Signal transformation and deep learning

The extracted raw signals were converted into 2D images using the Python PyTS package (27). We implemented a model training method in Python that employs Tensorflow, Keras, Scikit, Pandas, PyCM, and PyTS libraries (**Table S5**) (13, 25, 28–33). Keras implementations of ResNet-20 and ResNet-56 were slightly modified to support multi-gpu training, to adjust the learning rate scheduler, and to limit the channels to 1 and outputs to 4 classes (see Jupyter notebook in git repository v1.0.0 release source code). To drastically increase the speed of training, we employed Keras multi-GPU processing with Tensorflow-1.32. A Jupyter notebook presenting all commands used for the ResNet training protocol is available in the accompanying Github repository (release v1.0.0). Training was performed on a server with 4x NVIDIA V100 GPUs with 16GB memory each using NVLink.

## Data availability

Code, models, and scripts used demultiplex direct RNA reads—including example FAST5 data, data processing, reference sequences, and benchmarking scripts can be found at: https://github.com/Psy-Fer/deeplexicon. All FAST5 datasets used in this work will be made publicly available in SRA, under accession number PRJNA545820.

## Performance evaluation

ROC and precision metrics were performed using the ROCit package in R.

## Code availability

Code to demultiplex direct RNA reads, including example FAST5 data, can be found at: https://github.com/Psy-Fer/deeplexicon. All FAST5 datasets used in this work have been made publicly available in SRA, under accession number PRJNA545820.

## ACKNOWLEDGEMENTS

We would like to thank Dr. Tim Mercer for providing us with the sequin plasmids that have been used in this work. O.B. is supported by an international PhD fellowship (UIPA) from the University of New South Wales. MCL is supported by CRG International PhD Fellowships Programme. E.M.N was supported by a DECRA fellowship from the Australian Research Council (DE170100506) and is currently supported by CRG Severo Ochoa Funding. This work was funded by the Australian Research Council (DP180103571). We acknowledge the support of the Spanish Ministry of Economy, Industry and Competitiveness (MEIC) to the EMBL partnership, Centro de Excelencia Severo Ochoa and CERCA Programme / Generalitat de Catalunya.

## AUTHOR CONTRIBUTIONS

MAS, TE, JF and HL performed the bioinformatic analysis of the data and developed demultiplexing pipelines. TE designed and performed all deep learning models. MCL, OB and LB prepared the synthetic RNAs. MCL, OB, LB and KB prepared the direct RNA libraries and ran the sequencing. MAS and EMN conceived and supervised the project. MAS and EMN wrote the manuscript with assistance from all authors.

## REFERENCES

1. Pollard, M.O., Gurdasani, D., Mentzer, A.J., Porter, T. and Sandhu, M.S. (2018) Long reads: their purpose and place. Hum. Mol. Genet., 27, R234–R241.

2. Ardui, S., Ameur, A., Vermeesch, J.R. and Hestand, M.S. (2018) Single molecule real-time (SMRT) sequencing comes of age: applications and utilities for medical diagnostics. Nucleic Acids Res., 46, 2159–2168.

3. Rang, F.J., Kloosterman, W.P. and de Ridder, J. (2018) From squiggle to basepair: computational approaches for improving nanopore sequencing read accuracy. Genome Biol., 19, 90.

4. Garalde, D.R., Snell, E.A., Jachimowicz, D., Sipos, B., Lloyd, J.H., Bruce, M., Pantic, N., Admassu, T., James, P., Warland, A., et al. (2018) Highly parallel direct RNA sequencing on an array of nanopores. Nat. Methods, 15, 201–206.

5. Smith, A.M., Jain, M., Mulroney, L., Garalde, D.R. and Akeson, M. (2019) Reading canonical and modified nucleobases in 16S ribosomal RNA using nanopore native RNA sequencing. PLoS One, 14, e0216709.

6. He, K., Zhang, X., Ren, S. and Sun, J. (2016) Deep Residual Learning for Image Recognition. 2016 IEEE Conference on Computer Vision and Pattern Recognition (CVPR), 10.1109/cvpr.2016.90.

7. Hardwick, S.A., Chen, W.Y., Wong, T., Deveson, I.W., Blackburn, J., Andersen, S.B., Nielsen, L.K., Mattick, J.S. and Mercer, T.R. (2016) Spliced synthetic genes as internal controls in RNA sequencing experiments. Nat. Methods, 13, 792–798.

8. Liu, H., Begik, O., Lucas, M.C., Ramirez, J.M., Mason, C.E., Wiener, D., Schwartz, S., Mattick, J.S., Smith, M.A. and Novoa, E.M. (2019) Accurate detection of m6A RNA modifications in native RNA sequences. Nat. Commun., 10, 4079.

9. Ferguson, J.M. and Smith, M.A. SquiggleKit: A toolkit for manipulating nanopore signal data. Bioinformatics, 10.1093/bioinformatics/btz586.

10. Pak, M. and Kim, S. (2017) A review of deep learning in image recognition. In 2017 4th International Conference on Computer Applications and Information Processing Technology (CAIPT). pp. 1–3.

11. Eckmann, J.-P., -P Eckmann, J., Oliffson Kamphorst, S. and Ruelle, D. (1987) Recurrence Plots of Dynamical Systems. Europhysics Letters (EPL), 4, 973–977.

12. Wang, Z. and Oates, T. (2015) Encoding Time Series as Images for Visual Inspection and Classification Using Tiled Convolutional Neural Networks. In Workshops at the Twenty-Ninth AAAI Conference on Artificial Intelligence.

13. Gulli, A. and Pal, S. (2017) Deep Learning with Keras Packt Publishing Ltd.

14. van Dijk, E.L., Jaszczyszyn, Y., Naquin, D. and Thermes, C. (2018) The Third Revolution in Sequencing Technology. Trends Genet., 34, 666–681.

15. Leger, A., Amaral, P.P., Pandolfini, L. and Capitanchik, C. (2019) RNA modifications detection by comparative Nanopore direct RNA sequencing. BioRxiv.

16. Krause, M., Niazi, A.M., Labun, K., Torres Cleuren, Y.N., Müller, F.S. and Valen, E. (2019) tailfindr: alignment-free poly(A) length measurement for Oxford Nanopore RNA and DNA sequencing. RNA, 25, 1229–1241.

17. Workman, R.E., Tang, A.D., Tang, P.S., Jain, M. and Tyson, J.R. (2019) Nanopore native RNA sequencing of a human poly (A) transcriptome. Nature.

18. Ismail Fawaz, H., Forestier, G., Weber, J., Idoumghar, L. and Muller, P.-A. (2019) Deep learning for time series classification: a review. Data Min. Knowl. Discov., 33, 917–963.

19. Boža, V., Brejová, B. and Vinar, T. (2017) DeepNano: Deep recurrent neural networks for base calling in MinION nanopore reads. PLoS One, 12, e0178751.

20. Ni, P., Huang, N., Zhang, Z., Wang, D.-P., Liang, F., Miao, Y., Xiao, C.-L., Luo, F. and Wang, J. (2019) DeepSignal: detecting DNA methylation state from Nanopore sequencing reads using deep-learning. Bioinformatics, 10.1093/bioinformatics/btz276.

21. Teng, H., Cao, M.D., Hall, M.B., Duarte, T., Wang, S. and Coin, L.J.M. (2019) Chiron: translating nanopore raw signal directly into nucleotide sequence using deep learning (vol 7, giy037, 2018). Gigascience, 8.

22. Wick, R.R., Judd, L.M. and Holt, K.E. (2018) Deepbinner: Demultiplexing barcoded Oxford Nanopore reads with deep convolutional neural networks. PLoS Comput. Biol., 14, e1006583.

23. LeCun, Y., Bengio, Y. and Hinton, G. (2015) Deep learning. Nature, 521, 436–444.

24. Li, H. (2018) Minimap2: pairwise alignment for nucleotide sequences. Bioinformatics, 34, 3094–3100.

25. van der Walt, S., Colbert, S.C. and Varoquaux, G. (2011) The NumPy Array: A Structure for Efficient Numerical Computation. Computing in Science Engineering, 13, 22–30.

26. Savitzky, A. and Golay, M.J.E. (1964) Smoothing and Differentiation of Data by Simplified Least Squares Procedures. Analytical Chemistry, 36, 1627–1639.

27. Faouzi, J., Carryer, T., Lee, K.K., Yurchak, R. and Avis P (2019) johannfaouzi/pyts: Release of 0.7.3 version.

28. Abadi, M., Barham, P., Chen, J., Chen, Z., Davis, A., Dean, J., Devin, M., Ghemawat, S., Irving, G., Isard, M., et al. (2016) Tensorflow: A system for large-scale machine learning. In 12th ${USENIX} Symposium on Operating Systems Design and Implementation ({OSDI}$ 16). pp. 265–283.

29. Hunter, J.D. (2007) Matplotlib: A 2D Graphics Environment. Comput. Sci. Eng., 9, 90–95.

30. McKinney, W. and Others (2010) Data structures for statistical computing in python. In Proceedings of the 9th Python in Science Conference. Austin, TX, Vol. 445, pp. 51–56.

31. Faouzi, J. (2017) pyts: a Python package for time series transformation and classification.

32. Pedregosa, F., Varoquaux, G., Gramfort, A., Michel, V., Thirion, B., Grisel, O., Blondel, M., Prettenhofer, P., Weiss, R., Dubourg, V., et al. (2011) Scikit-learn: Machine Learning in Python. J. Mach. Learn. Res., 12, 2825–2830.

33. Haghighi, S., Jasemi, M., Hessabi, S. and Zolanvari, A. (2018) PyCM: Multiclass confusion matrix library in Python. JOSS, 3, 729.

