## Supplementary material for "Barcoding and demultiplexing Oxford Nanopore native RNA sequencing reads with deep residual learning": Supplementary_info.pdf

**SUPPLEMENTARY FIGURES**

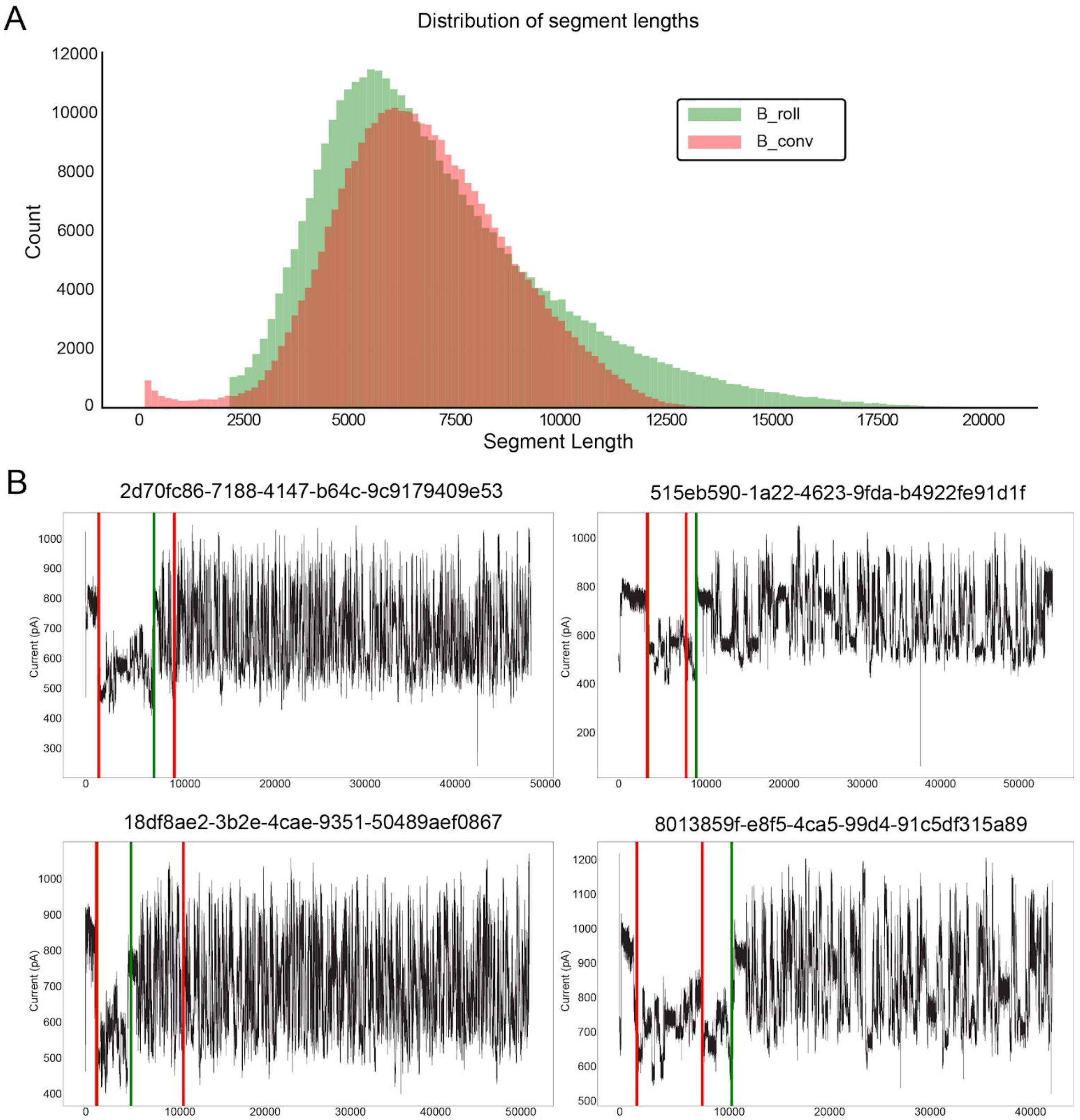


**Figure S1.** Comparison of barcode segmentation strategies. **(A)** Distribution of extracted signal lengths (red = *B_conv*, green = *B_roll*). **(B)** Example of barcode segmentation positions in signal data from four reads using the two tested segmentation algorithms (red = *B_conv*, green = *B_roll*).

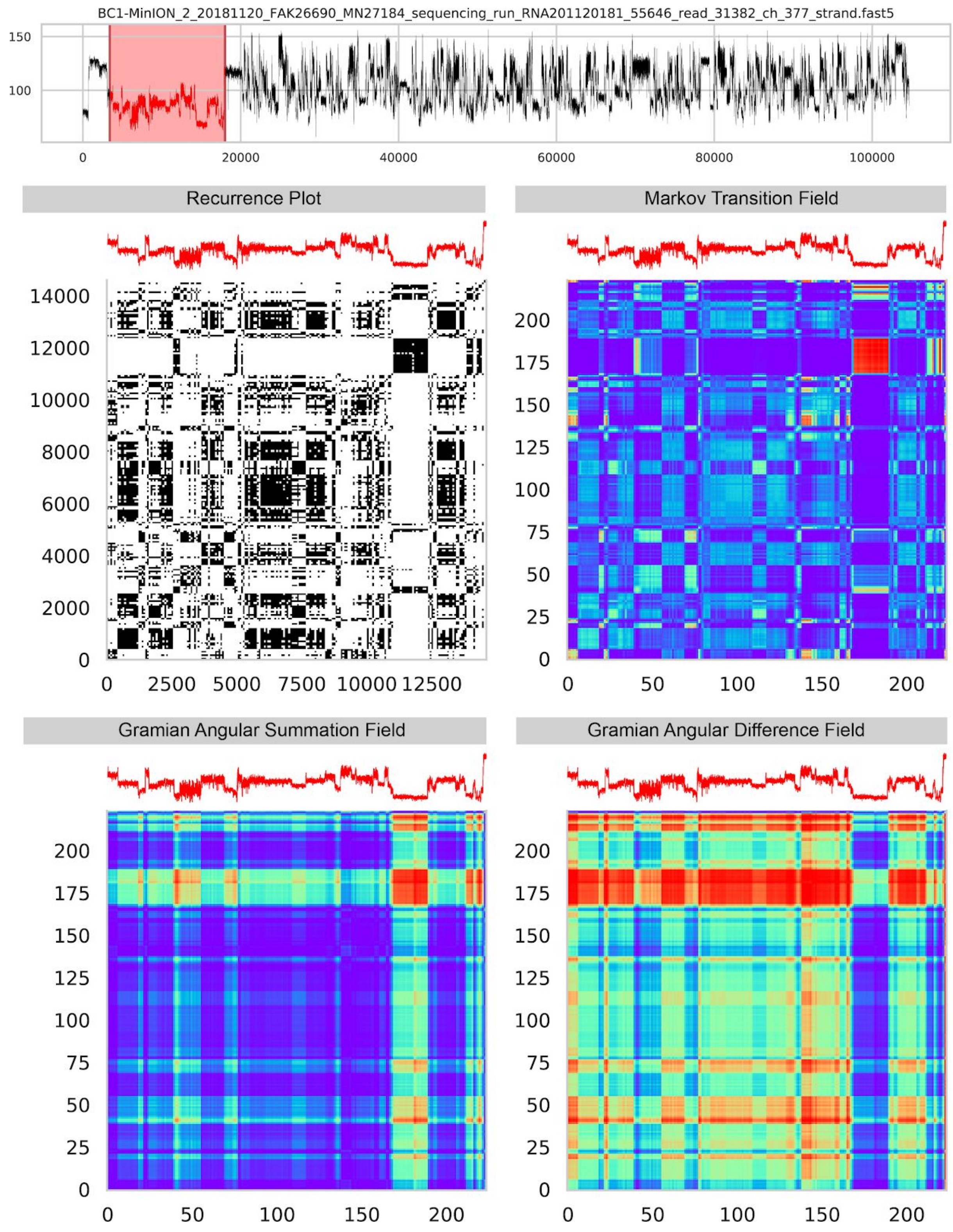


**Figure S2. Transformation of adapter raw signals into 2D images.** Example illustrating how raw signal is converted into **(top)** Recurrence Plot and Markov Transition Field, and **(bottom)** Gramian Angular Summation (GASF) and Gramian Angular Difference (GADF)

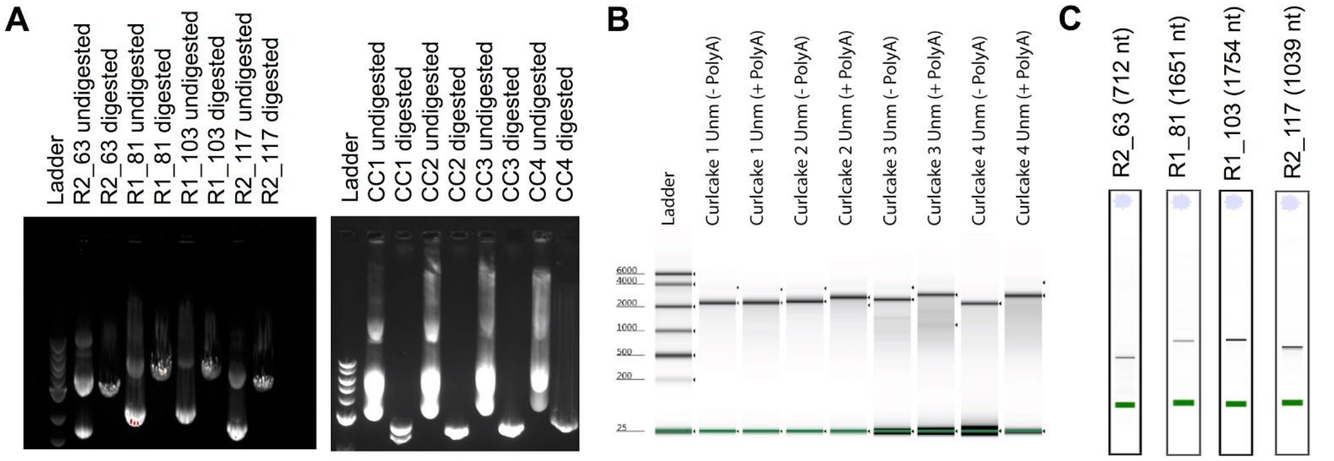


**Figure S3. Quality control of the production of *in vitro* transcribed sequences, which were ligated to custom barcoded adapters. (A)** Sequin and Curlcake plasmid digestion was confirmed by agarose gel (**B,C**) Correct IVT product lengths were confirmed using Bioanalyzer -in the case of sequins- (B) ,or TapeStation -in the case of curlcakes- (C).

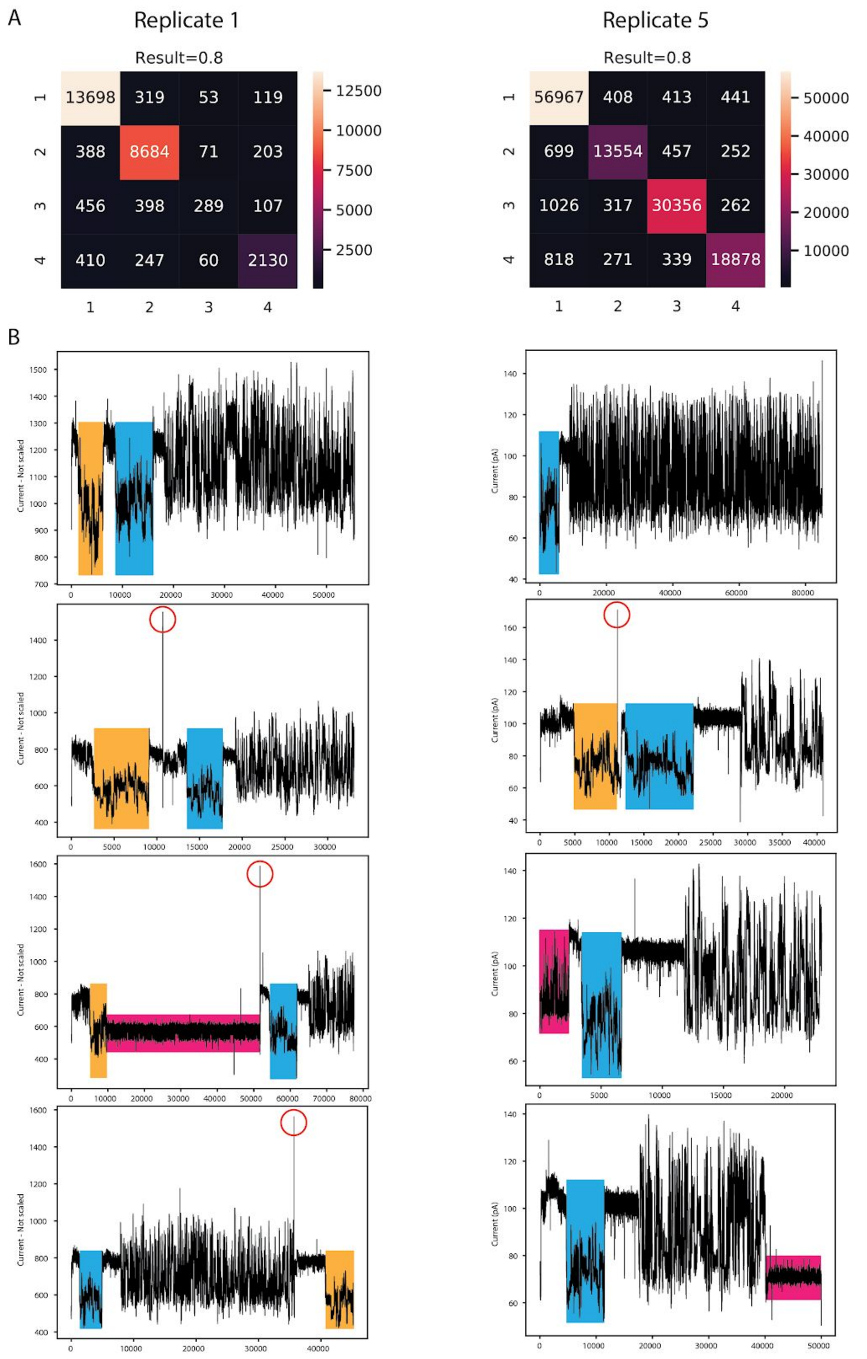


**Figure S4. Performance of Deeplexicon on independent replicates (replicate 1 and replicate 5) not used for training or testing. (A)** Confusion matrices for replicate 1 (left) and replicate 5 SQK-RNA002 (right). Replicate 1 was prepared using SQK-RNA001 library preparation kit and was sequenced in a R9.5 flowcell, whereas replicate 5 was prepared using SQK-RNA001 library preparation kit and sequenced in a R9.4.1 flowcell. **(B)** Examples illustrating chimeric artefacts during library preparation and/or errors in MinKNOW read assignment, which were most frequent in replicate 1. These artefacts impact the ability to correctly identify the adapter. The correct adapter/barcode within the RNA read is highlighted in blue; a second adapter/barcode detected within the read is shown in orange, and should belong to a different read; signal artefacts, which may affect the identification of the barcoded region, are highlighted in pink. Red circles depict a very short “open pore” state that MinKNOW software missed, and therefore failed to cut the signal into two indendent reads.

### SUPPLEMENTARY TABLE LEGENDS

**Table S1.** Barcode sequences and custom oligonucleotides (oligoA and oligoB) employed to prepare barcoded direct RNA sequencing libraries.

**Table S2.** Sequencing metrics for the three barcoded runs used to train, test and validate the demultiplexing algorithm

**Table S3.** Quality analysis of *in vitro* transcribed RNA products and library preparation

**Table S4**. Reference fasta sequences of the 4 Sequins and 4 Curlcakes used in this work
